# Microbial community composition of translocated ancient woodland soil: a case study

**DOI:** 10.1101/2021.06.30.450580

**Authors:** Nicolas Borchers, Jacqueline Hannam, Mark Pawlett

## Abstract

Soil translocation is an ecological habitat restoration technique which consists of moving the entire topsoil from a donor site to a chosen receptor site. We investigated changes in soil chemistry and microbiology three years following the salvage of semi-ancient woodland soil and materials (0.94 ha) to a nearby receptor pasture due to road widening works (Kent, UK). We sampled i) intact woodland soils adjacent to the area of soils that was translocated to represent the lost donor site, ii) the soil three years after it had been translocated, and iii) grassland soils adjacent to the translocated soil to represent the original receptor site. The intention was to ascertain if shifts in soil chemistry and microbial community composition (Phospholipid Fatty-acid analysis: PLFA) occurred due to soil translocation. PLFA signature biomarkers demonstrated the overall microbial community profile of the translocated and woodland soils were similar; however, salvaged soils had a 40% increase in the Arbuscular Mycorrhizal Fungi (AMF) bioindicator fatty acid 16:1ω5, a 10% decrease in the Gram-positive bacterial fatty acids, and increased pH (5.01-5.77) compared to the original donor woodland soil. The AMF bioindicator and the first Principal Component (PC1) of the PCA of PLFA data positively correlated with soil pH (*r^2^*=0.94 and *r^2^*=0.88 respectively) across all three experimental groups. Considering that soil pH increases with depth in this location, it is likely that mixing of soil horizons during translocation increased the topsoil pH causing changes in the soil microbial communities. We concluded that after three years, the chemical and microbial properties of the salvaged soil were characteristic of a woodland soil but showed signs of disturbance.

**Implications for practice:**

- Translocated semi-ancient woodland soils may still exhibit similar microbiological characteristics to the donor site three years after being moved to a pasture receptor site.
- However, disturbances expressed as significant changes in chemistry and microbial communities can affect soils following their translocation.
- Where soil pH varies with depth, translocation by loose-tipping may redistribute acidity within the soil profile, with important cascading impacts on the soil microbial communities.

## Introduction

Soil translocation is a recent, last-resort and expensive technique to salvage or restore ecosystems. The technique entails removing the topsoil from a donor site, which typically would otherwise be lost to development, and its transfer to a chosen receptor site, where a similar ecosystem is to be created or restored (Anderson 2003). Compared to restorations based solely on the planting of target vegetation communities, the aim of soil translocation is to eventually restore or create an ecosystem in a state closer to the original donor, and to attain this target in a shorter amount of time (Anderson 2003; Ryan 2013). Translocated soil as a medium not only provides a similar physical and chemical soil environment, but also a similar biological environment to the donor site since the seed bank, soil macro- and meso-fauna, and soil microbes are transferred in the process. Soil microbial communities have adapted specifically to different ecosystems and land uses, and these microbes together with the network of associations they form with plants are essential to the whole above- and below-ground ecosystem function (Wardle et al. 2004; Morris et al. 2006; Van Der Heijden et al. 2008). In particular, mycorrhizal plant-fungi associations have a large influence on vegetation growth and community assemblage. Some of those associations are very specific (Helgason et al. 2007) and are of practical relevance to ecosystem restoration projects (Hoeksema et al. 2010).

There are few available cases studies of salvage translocation of ancient woodland soil and materials, either because it has rarely been attempted, because monitoring is lacking or studies are not published (Ryan 2013), or because ancient woodland has protected status so alternative routes are probably favored in the planning process. Most of the published research has evaluated the translocation of herbaceous environment such as grassland, wetland or heathland, whose soil and vegetation can more easily be removed by machinery compared to woodland (Bullock 1998; Humphries 2010). The few studies available on the salvage translocation of temperate woodland mostly focused on above-ground vegetation as indicators of the restoration trajectory, either in situ (Helliwell et al. 1996; Hietalahti & Buckley 2000; Craig et al. 2015; Buckley et al. 2017), or ex situ (Craig & Buckley 2013). Only one identified study turned its attention to soil chemistry and soil respiration. In a one-year field experiment immediately following the translocation of deciduous woodland soil (Kent, UK), Hietalahti et al. (2005) measured on a monthly basis soil respiration, mineral nitrogen and organic matter but found no evidence of increased carbon and nitrogen mineralisation rates.

Microbial communities, in addition to their role as facilitators in the restoration process, could also behave as followers of ecosystem changes, and therefore serve as valuable indicators to evaluate the progress and guide the management of ecological restoration projects. In addition, characterizing the soil microbes can be used to conclusively distinguish between systems and ecological trajectories, as well as detect disturbances and degradation (Harris 2003, 2009). Studies have successfully used complementary analysis of community structure through PLFA analysis combined with functional profiles to investigate the transformations in soil microbial communities in the context of ecosystem restoration or perturbation (Zogg et al. 1997; Kourtev et al. 2003; Andersen et al. 2010). This approach was followed by Case (2017, unpublished) in the only identified study investigating the effects of woodland soil translocation concurrently on soil chemistry as well as on microbial function and structure using PLFA biomarkers. This research compared a salvaged ancient woodland soil in Kent, UK to soil from an adjacent undisturbed woodland eighteen years following the translocation. It found that soil chemistry was essentially similar but that soil microbes experienced a shift from bacterial dominated communities in the donor woodland soil to fungal-dominated communities in the translocated soil.

Here we investigated the changes in chemistry (pH, soil organic matter, available nitrogen and phosphorus) and microbiology (microbial biomass, activity and PLFA) of a translocated woodland soil in Kent, UK, three years after translocation, compared to adjacent original woodland soil. Our hypothesis was that the translocated soil would still show characteristics of a woodland soil three years after translocation.

## Methodology

### Study site

The study area (Fig. 1) is located along the eastern side the A21 road between Tonbridge and Pembury, Kent, England (51° 10.417’ N, 0° 18.035’ E, altitude 116m). The climate is temperate oceanic with average annual max/min temperatures of 14.2/6.9°C and 833 mm annual precipitation (MET Office). The soils are Eutric Luvic Planosols. These soils are seasonally waterlogged and are fine silty, fine loamy or clayey, above clayey subsoil (National Soil Resources Institute UK). Due to widening works on the A21, the topsoils from a woodland measuring 0.94 ha (“donor site”) were salvaged and translocated in 2015 to a nearby pasture (“receptor site”). The floral ecology of the donor site was surveyed by Smith (2014) and assessed as “semi-natural ancient woodland” corresponding in the UK National Vegetation Classification (Hall et al. 2001) to a W10c community (*Quercus robur* - *Pteridium aquilinum* - *Rubus fruticosus*, sub-community *Hedera helix*) with significant patches of W8d (*Fraxinus excelsior* - *Acer campestre* - *Mercurialis perennis*, sub-community *Hedera helix*). The receptor site was a semi-improved grassland (i.e. species poor, dominated by *Holcus lanatus* and *Festuca rubra* with frequent *Trifolium repens)*, used as pasture since at least 1940. The donor site was felled and cleared in early 2015, and in March 2015 its topsoil was removed using excavators equipped with toothless buckets. Insofar as practically possible, the soil was scraped down to the visible subsoil boundary, and in any case to a minimum of 10cm depth. Subsoil was therefore potentially removed along with the topsoil. The donor soils were then translocated with dumpers to the prepared receptor site, which had previously been stripped of its topsoil to a depth of 30cm, and were loose-tipped and spread evenly. Selected coppice stools and small saplings were also moved from donor to receptor, supplemented after translocation by the planting of saplings of local provenance stock (Smith 2014).

**Figure 1:**
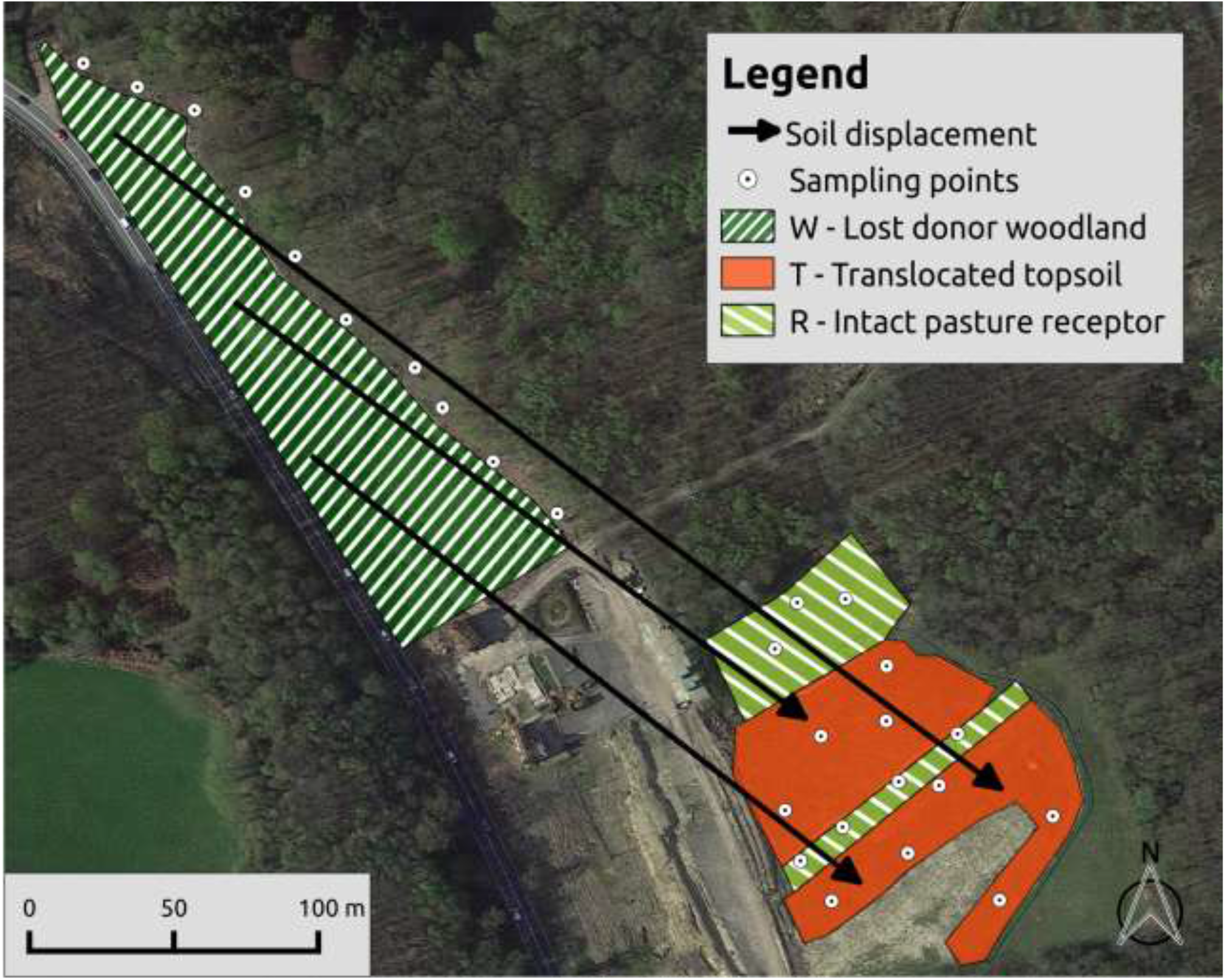
Map of the study site

### Experimental design

Topsoil was sampled on 31 October 2017 from three different locations (Fig. 1). These were: (1) A reference woodland area chosen as a proxy for the lost donor site (group “W”, n=10). Sampling was done within a band immediately adjacent to the donor site to ensure similar common vegetation and ecological history to the donor (Smith 2014). Sampling depth was consistent with the topsoil that was removed from the donor site, i.e., down to the visible topsoil/subsoil boundary but to a minimum of 10 cm depth, even if this included subsoil.

(2) Translocated soil (group “T”, n=9), which was sampled down to the clearly visible boundary with the underlying original receptor subsoil.

(3) A reference intact receptor site (group “R”, n=7). Samples were taken from the areas of the pasture left untouched, to a depth of 15 cm.

At each sampling point 4 to 7 soil cores were obtained using a Dutch auger, and bulked as a composite sample, then fresh sieved at 2mm. Sample point locations were taken at a minimum distance of two meters from any mature tree. Three subsoils, one per group, were also sampled for pH only. Samples were kept refrigerated and analysed later for available nitrogen (N), microbial biomass, and microbial activity, frozen at −85°C then freeze-dried for PLFA, and air dried for pH, organic matter (OM) and available phosphorus (P).

### Chemical analysis

Soil pH was determined on air-dried samples in a water extract (1:5 v/v), according to BS ISO 10390:2005. The OM content was determined by measuring the loss of mass after ashing air-dried soil at 450°C for 4 hours, according to BS EN 13039:2000. Ammonium-N, nitrate-N and nitrite-N were determined on fresh soil by extraction in a 2M potassium chloride solution and measuring the respective concentrations by spectrometry, according to method 53 of the MAFF Reference Book RB427 (MAAF 1986). The reported available N value is the sum of ammonium-N, nitrate-N and nitrite-N. Available phosphorus was measured on air-dried soil by extraction in a 0.5M sodium hydrogen carbonate solution (Olsen’s P) and measuring the phosphorus concentration by spectrometry, according to BS: 7755: Section 3.6:1995 (ISO 11263:1994).

### Microbial analysis

Microbial biomass-C was determined on fresh soil by fumigation-extraction, according to Jenkinson and Powlson (1976). The microbial activity was calculated as the arithmetic average of triplicate measurements using the Rapid Automated Bacterial Impedance Technique (RABIT®) device (Don Whitley Scientific Ltd) and following the methodology described in Ritz et al. (2006). The relative abundances (% mol) of 34 PLFA were measured following a method adapted from Frostegård et al. (1991), as described in detail in Courtney et al. (2014). Bioindicators fatty acids used included: the sum of i15:0, ai15:0, i16:0, i17:0, ai17:0, 10me18:0 - Gram-positive (G^+^) bacteria (Zelles 1999), the sum of 16:1ω7c, 17:1ω7, 18:1ω7t, cyc17:0, cyc19:0 - Gram-negative (G^−^) bacteria (Zelles 1999), 18:2ω6,9 - ectomycorrhizal and saprophytic fungi (Frostegård & Bååth 1996; Frostegård et al. 2011), 16:1ω5 - Arbuscular Mycorrhizal Fungi (AMF) (Olsson 1999; Frostegård et al. 2011). Total bacteria were calculated as the sum of G^+^ and G^−^ bacteria. A fungal: bacterial ratio was calculated as 18:2ω6,9 divided by total bacteria (Bååth & Anderson 2003).

### Statistical analysis

Statistical analyses were carried out in R 4.0.3. Data were first tested for normality and homoscedasticity. OM was log-transformed for statistical analysis. Soil pH, OM, available N, available P, microbial biomass, microbial activity and the PLFA bioindicators were each subjected to a one-way analysis of variance (ANOVA), followed by post hoc Tukey-Kramer multi-comparison tests. The relative abundance (% mol) of the 34 identified fatty acids were subjected to Principal Component Analysis (PCA, correlation matrix based, package *vegan*) as well as permutational multivariate analysis of variance (PERMANOVA) using *adonis* in package *vegan*. The post hoc pairwise comparison was performed using the package *pairwiseAdonis* and adjusted with Bonferroni correction. Relationships between PLFA bioindicators, PCA principal components and soil chemistry were investigated using stepwise multiple linear regression analysis with BIC (Bayesian information criterion) as selection criterion (package *stats*). All differences were considered statistically significant if *p*<0.05.

## Results

### Chemical parameters

Soil OM, available N and available P were not significantly different between the donor woodland and translocated soil (Table 1), however pH showed an increase from 5.01 to 5.77 (*p*<0.05). The pasture receptor soil had a higher pH (5.93 to 5.01: *p*<0.05) compared to the woodland donor soil and greater N availability (8.23 to 3.58 μg-N/g: *p*<0.05) compared to the translocated soil. OM and available P were not statistically different (*p*>0.05) between the different areas. The three subsoils had a pH of 4.46 (W), 6.49 (T) and 7.52 (R), to be compared to corresponding topsoil pH at the same locations of 4.34, 5.59 and 5.98 respectively.

**Table 1:**
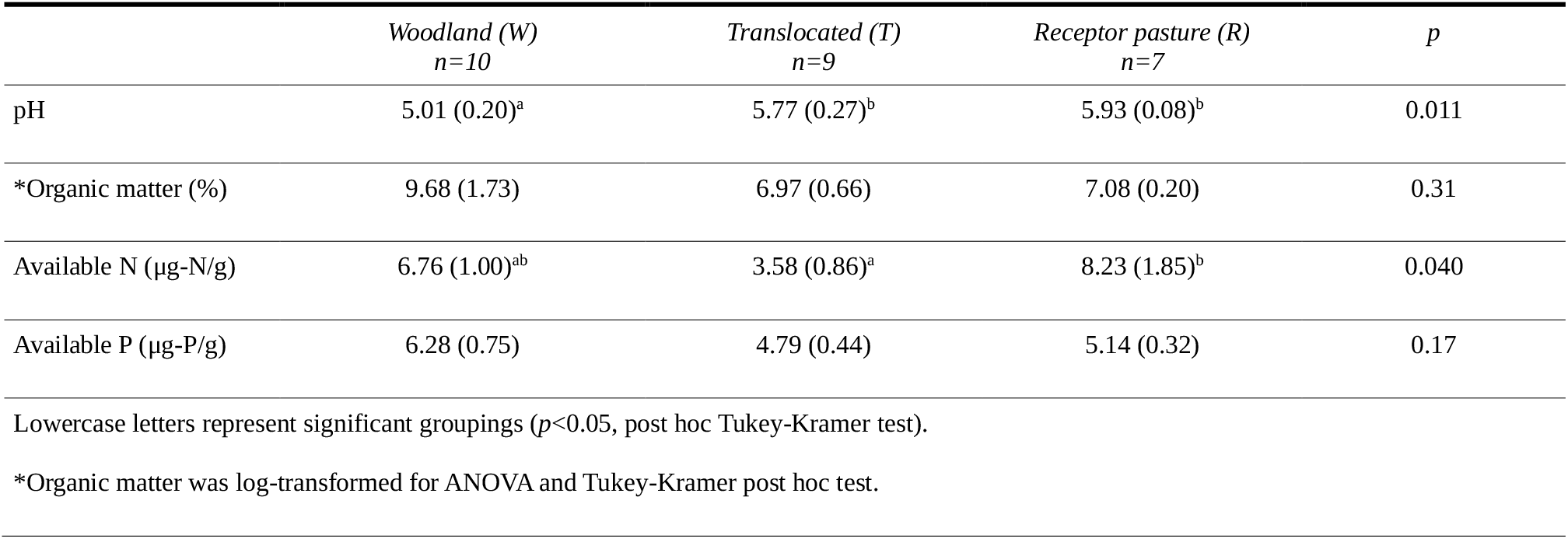
Soil chemical indicators. Values are means with SE in parenthesis.

|  | Woodland (W)<br>*n*=10 | Translocated (T)<br>*n*=9 | Receptor pasture (R)<br>*n*=7 | *p* |
| --- | --- | --- | --- | --- |
| pH | 5.01 (0.20)<sup>a</sup> | 5.77 (0.27)<sup>b</sup> | 5.93 (0.08)<sup>b</sup> | 0.011 |
| *Organic matter (%) | 9.68 (1.73) | 6.97 (0.66) | 7.08 (0.20) | 0.31 |
| Available N ($\mu\text{g-N/g}$) | 6.76 (1.00)<sup>ab</sup> | 3.58 (0.86)<sup>a</sup> | 8.23 (1.85)<sup>b</sup> | 0.040 |
| Available P ($\mu\text{g-P/g}$) | 6.28 (0.75) | 4.79 (0.44) | 5.14 (0.32) | 0.17 |

Lowercase letters represent significant groupings ( $p<0.05$ , post hoc Tukey-Kramer test).
\*Organic matter was log-transformed for ANOVA and Tukey-Kramer post hoc test.

### Microbial parameters

The majority of microbial indicators (biomass, activity, fungi as 18:2ɷ6, 9, the total and G^−^ bacterial fatty acids, and the fungal:bacterial ratio) were not significantly different between the donor woodland and translocated soil (Table 2). However the AMF bioindicator (16:1ω5) showed an increase (2.7 to 3.7% mol: *p*<0.05), and the G^+^ bacterial bioindicator a decrease (16.9 to 15.3% mol: *p*<0.05) in the translocated soil compared to the woodland donor. The pasture receptor soil had reduced fungi (4.2 to 7.5% mol: *p*<0.05), greater AMF (4.4 to 2.7% mol: *p*<0.01), and reduced fungal:bacterial ratio (0.08 to 0.16: *p*<0.05) compared to the woodland donor soil; and reduced fungi (4.2 to 7.3% mol: *p*<0.05), greater G^+^ (17.2 to 15.3% mol: *p*<0.05), and reduced fungal:bacterial ratio (0.08 to 0.16: *p*<0.05) compared to the translocated soil. The microbial biomass, microbial activity, G^−^ and total bacterial bioindicators were not statistically different (*p*>0.05) between the different groups.

**Table 2:** Soil microbiological indicators. Values are means with SE in parentheses.

|  | Woodland (W)<br>*n*=10 | Translocated (T)<br>*n*=9 | Receptor pasture (R)<br>*n*=7 | *p* |
| --- | --- | --- | --- | --- |
| Microbial biomass (µg-C/g) | 502 (97) | 501 (70) | 733 (47) | 0.11 |
| Microbial activity (µg-C g<sup>-1</sup> h<sup>-1</sup>) | 1.54 (0.14) | 1.66 (0.23) | 1.39 (0.14) | 0.58 |
| Fungi (% mol) | 7.5 (0.9)<sup>a</sup> | 7.3 (0.4)<sup>a</sup> | 4.2 (0.5)<sup>b</sup> | 0.008 |
| AMF (% mol) | 2.7 (0.3)<sup>a</sup> | 3.7 (0.4)<sup>b</sup> | 4.4 (0.1)<sup>b</sup> | 0.003 |
| Bacteria (% mol), of which: | 48.7 (0.8) | 47.1 (0.7) | 49.7 (0.6) | 0.074 |
| Gram-positive (% mol) | 16.9 (0.6)<sup>a</sup> | 15.3 (0.4)<sup>b</sup> | 17.2 (0.3)<sup>a</sup> | 0.020 |
| Gram-negative (% mol) | 31.8 (0.9) | 31.8 (0.6) | 32.5 (0.5) | 0.75 |
| Fungal:bacterial ratio | 0.16 (0.02)<sup>a</sup> | 0.16 (0.01)<sup>a</sup> | 0.08 (0.01)<sup>b</sup> | 0.019 |
| Lowercase letters represent significant groupings ($p<0.05$, post hoc Tukey-Kramer test). | | | | |

The first two principal components (PC) of the PCA of PLFA data cumulatively accounted for 58.6% of the total variation, with PC1 and PC2 accounting for 43.6% and 15.0% respectively (Fig. 2). Fatty acids with a positive loading (>0.8) on PC1 included: ai15:0 (G^+^), 16:1ω11t, 16:1ω5 (AMF) and 18:1ω7t (G^−^). Fatty acids with a negative loading (<−0.8) on PC1 included: i15:0 (G^+^), i16:0 (G^+^) and UK 42.41 (Fig. 2). Fatty acids did not correlate to PC2 (>0.8 or <−0.8). The separation of the centroids of the distributions of the three groups was statistically significant (PERMANOVA, *p*<0.001). The receptor pasture soil differed significantly from both the donor and the translocated soil in the separation of their centroids (*p*<0.05, post hoc PERMANOVA pairwise comparison adjusted with Bonferroni correction), however the donor and translocated soil did not differ significantly (*p*=0.12).

**Figure 2:**
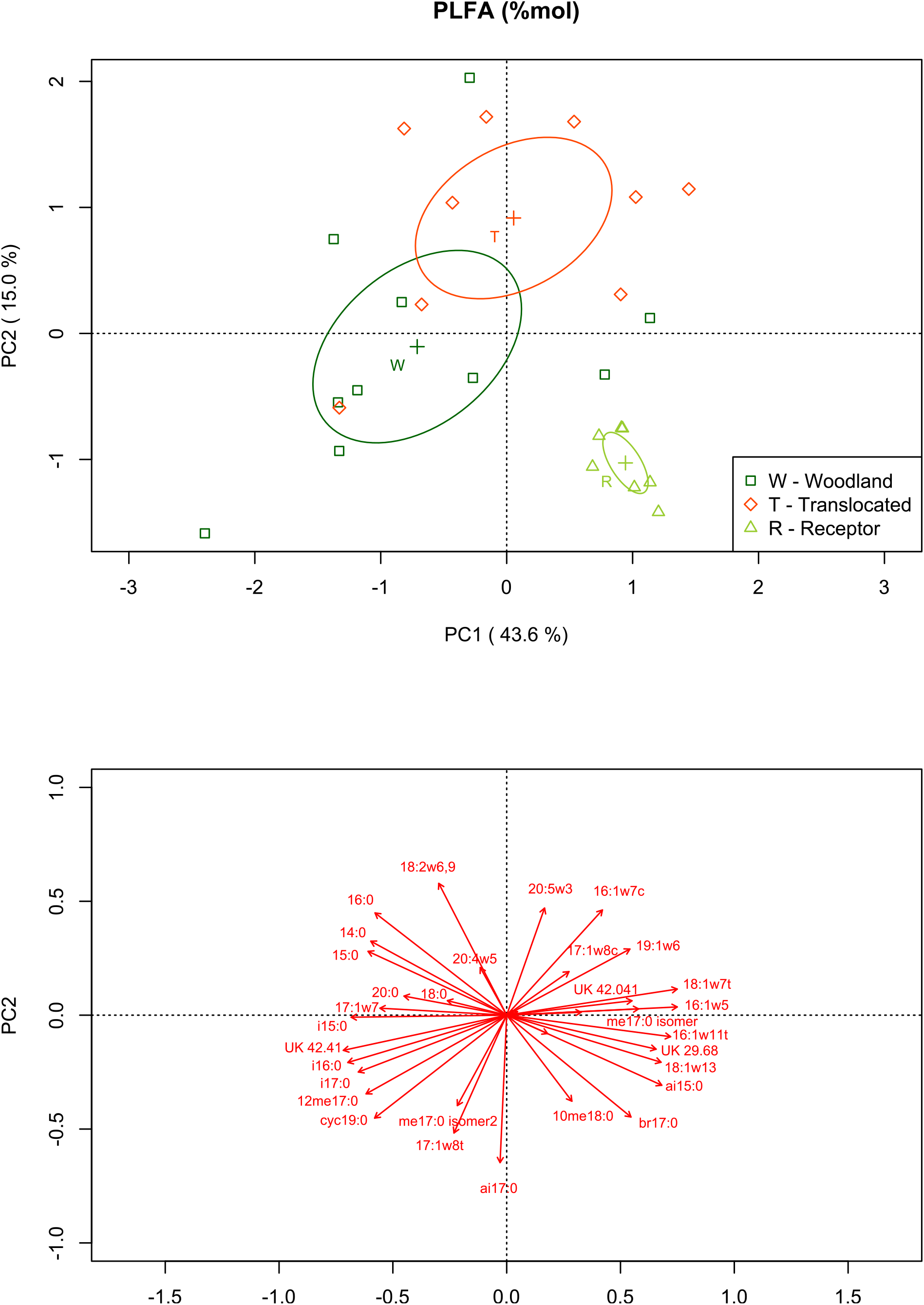
PLFA - Principal Component Analysis. PCA scores (top) and loadings (bottom). Ellipses on scores plot represent 95% confidence intervals of the mean samples for W, T and R respectively.

Multiple stepwise regression using the BIC criterion applied to the PLFA bioindicators AMF, fungi, G^+^ and G^−^ as well as to the first two PCA principal components, using the soil chemistry indicators (pH, OM, available N and available P) as explanatory variables, indicated that the variability in G^+^ and G^−^ bacteria can only be weakly explained by soil chemistry (*r^2^*=0.26, *p*=0.008; *r^2^*=0.29, *p*=0.004, respectively), while no satisfactory model could be fitted to fungi. AMF and PC1 however correlated well with soil chemistry (*r^2^*=0.94, *p*<0.001; *r^2^*=0.88, *p*<0.001, respectively), the best fit models retaining pH as sole explanatory variable (Fig. 3). No satisfactory model could be fitted to PC2. The strong correlations between PC1 and pH on one hand, and AMF and pH on the other hand, are linked due to the important contribution of the AMF bioindicator 16:1ω5 to PC1 (loading = 0.88, Fig. 2).

**Figure 3:**
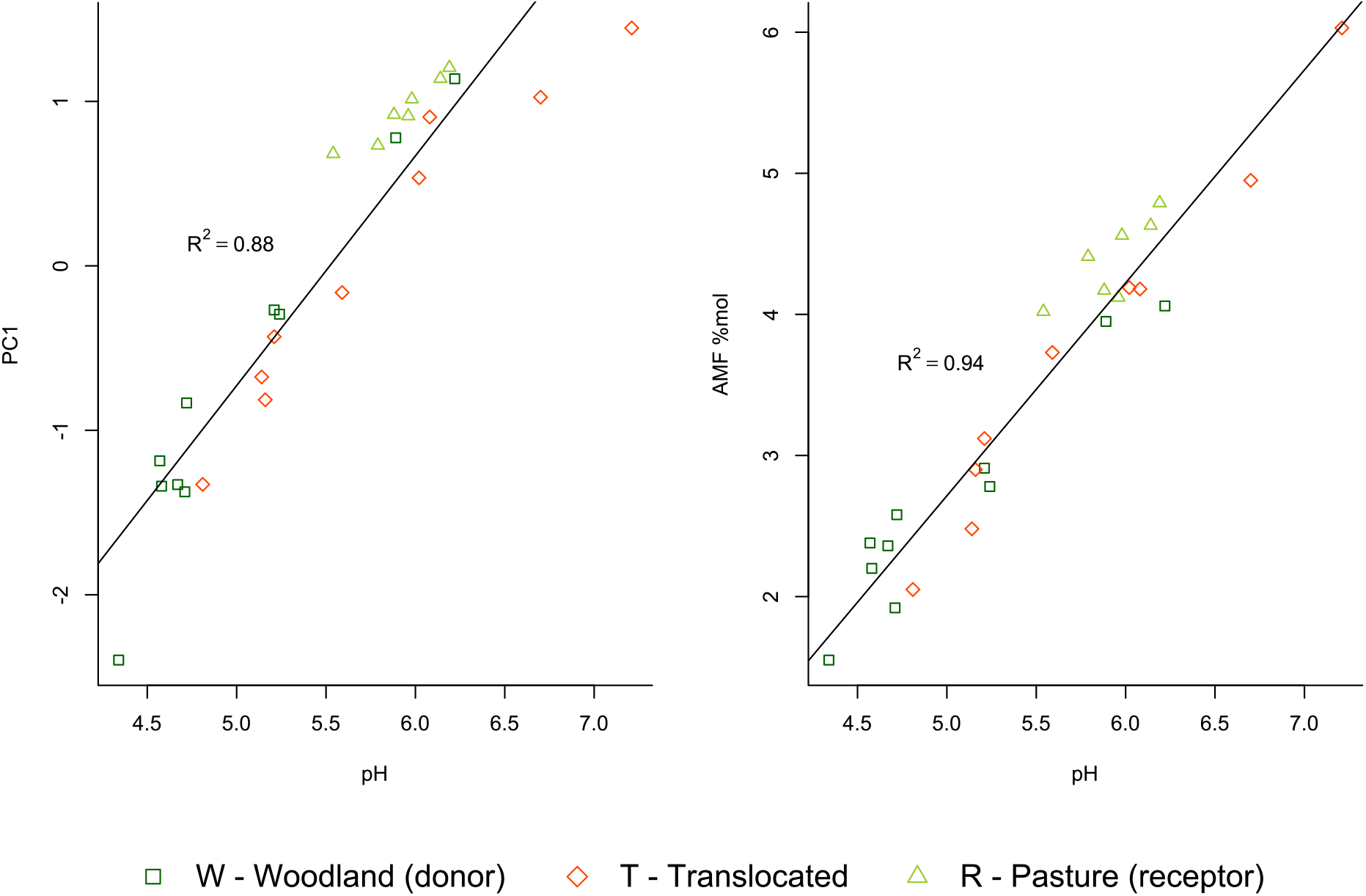
Amongst all microbial indicators, PC1 and the AMF bioindicator were best correlated with soil chemistry. Stepwise multiple regression using all four measured soil chemical parameters retained pH as sole explanatory variable in the best fit models.

## Discussion

Three years following translocation, overall microbiological indicators suggest that the translocated soil did not differ significantly compared to the original woodland soil. However, there was an increase (+40%) in the AMF and a decrease (−10%) in the G^+^ bacterial fatty acids. The only significant change in chemical parameters was an increase in pH (5.01 to 5.77). These findings are in good agreement with Case (2017). Using an analogous methodology on a similar site in Kent, UK, eighteen years following translocation they reported in the translocated soil a significant increase in pH (4.63 to 5.44) and AMF bioindicator (+50%) as well as decreases in the G^+^ bacterial bioindicator (−18%) compared to donor woodland soil. They also measured increases in the fungal bioindicator (+126%) and the G^−^ bacterial bioindicator (+19%) in the translocated soil, which were not observed here.

Soil pH is a major factor structuring soil microbial communities (Rousk, Bååth, et al. 2010). We found a robust correlation between pH and the PLFA profile, especially the 16:1ω5 AMF bioindicator, closely reflecting the findings of other studies in arable and forest soils (Bååth & Anderson 2003; Rousk, Brookes, et al. 2010). The same studies have however shown that the large change in abundance of certain fatty acids (in particular 16:1ω5) along pH gradients only partly represents an actual change in microbial community composition, and may in large parts be due to other pH related effects, resulting in an overestimation of the correlation. Furthermore, associating PLFA to specific microbial groups should be done with caution, since many are found across a range of organisms, in particular 16:1ω5 which is found both in AMF and in bacteria (Frostegård et al. 2011). Despite these risks of partial erroneous attribution, our results suggest that pH is the main driver in the observed shifts in the microbial communities. These microbial changes may also indicate an initial disturbance response from the translocation of soil, possibly followed by a partial recovery.

Case (2017) postulates that the observed increase in soil pH may be due to mixing of the woodland topsoil with less acidic subsoil at the donor site during removal. Soil acidity in woodland generally decreases with depth (Blake et al. 1999), as was observed on our study between the topsoil and subsoil. Consequently, differences in measured pH values, and possibly in other depth-dependent parameters between donor and translocated soil, could be an artifact of sampling depth rather than reflect effective changes in these properties. While we ensured by visual inspection that no receptor subsoil was included while sampling the translocated soil, it is still uncertain whether the sampling depth on the donor site was representative of the soil thickness effectively removed and translocated. Nevertheless, it is highly probable that handling the soil redistributed acidity within the soil profile during the translocation.

Other factors affecting the translocated soil could be due to interactions with the surrounding receptor soil. Compared to the donor woodland, the topsoil of the receptor site under pasture was significantly less acidic and had lower fungi, higher AMF, lower fungal:bacterial ratio and a significantly different PLFA profile (PERMANOVA, *p*<0.05). Such dissimilarities in donor and receptor soils generally justify decisions to remove the receptor topsoil prior to translocating the donor soil (Anderson 2003), as was carried out here. Even with prior topsoil removal, the dissimilarities between donor and receptor could affect the translocated soil through horizontal and vertical boundary effects including direct mixing during translocation, subsequent bioturbation, chemical exchanges, and biological inoculation. The different environmental conditions at the receptor site (early successional vegetation) compared to the donor site (mature woodland) may be an equally important driver of changes affecting the soil, in particular due to differences in plant-soil interactions and shading (Helliwell et al. 1996; Anderson 2003; Hietalahti et al. 2005; Ryan 2013; Craig et al. 2015). Specifically, decreased litter deposition at the receptor site may reduce acidic inputs and production in the upper soil horizons (Blake et al. 1999). However, our results do not suggest that after three years the translocated soil is on an intermediate trajectory between the original woodland soil and the receptor pasture soil, as the various soil parameters do not uniformly point in one direction of change towards or away from pasture soil.

In conclusion, we postulated that three years after being translocated, the salvaged soil would still show characteristics of a woodland soil. We found that the microbial phenotypical profile of the translocated soil was overall not significantly different from the donor soil, but there was an increase in pH and AMF and a decrease in gram-positive bacteria, indicative of disturbance. Soil pH is likely the main driver behind the observed microbial changes. The pH increase could tentatively be linked to the mixing of the donor soil profile with more neutral subsoil during the salvage operation. Changes in soil microbiology could furthermore be due to the influence of the surrounding receptor soil matrix, or to the different vegetation and shading at the receptor site. The knowledge gained will be valuable to assess the suitability of soil translocation as a technique for future woodland salvage and restoration projects.

## Acknowledgments

We thank Bruce Lascelles from Arcadis for providing the opportunity and funding for the field and lab work of the study, as well as Miguel Rodrigo Matos Correra Pereira Nunes for his help with the collection of soil samples. Technical assistance was provided by Alan Nelson, Ceri Dawson, Ian Truckell, Nuannat Simmons, Paul Barton and Caitlin Hedinburgh-Bailey.

Data underlying this study is openly available through the Cranfield University repository at https://doi.org/10.17862/cranfield.rd.14882727

## Notes

https://doi.org/10.17862/cranfield.rd.14882727

### Competing Interest Statement

The authors have declared no competing interest.

## LITERATURE CITED

Andersen R, Grasset L, Thormann MN, Rochefort L, Francez AJ (2010) Changes in microbial community structure and function following Sphagnum peatland restoration. Soil Biology and Biochemistry 42:291–301

Anderson P (2003) Habitat translocation - a best practice guide. CIRIA, London.

Bååth E, Anderson TH (2003) Comparison of soil fungal/bacterial ratios in a pH gradient using physiological and PLFA-based techniques. Soil Biology and Biochemistry 35:955–963

Blake L, Goulding KWT, Mott CJB, Johnston AE (1999) Changes in soil chemistry accompanying acidification over more than 100 years under woodland and grass at Rothamsted Experimental Station, UK. European Journal of Soil Science 50:401–412

Buckley P, Helliwell DR, Milne S, Howell R (2017) Twenty-five years on – vegetation succession on a translocated ancient woodland soil at Biggins Wood, Kent, UK. Forestry: An International Journal of Forest Research 90:561–572

Bullock JM (1998) Community translocation in Britain: Setting objectives and measuring consequences. Biological Conservation 84:199–214

Case E (2017) Soil characteristics of a woodland created using translocated, loose-tipped, ancient woodland topsoil in Kent, England. MSc Thesis. Cranfield University, UK

Courtney R, Harris JA, Pawlett M (2014) Microbial Community Composition in a Rehabilitated Bauxite Residue Disposal Area : A Case Study for Improving Microbial Community Composition. 22:798–805

Craig M, Buckley GP (2013) Responses of two woodland geophytes to disturbance caused by soil translocation. Plant Ecology 214:1091–1103

Craig M, Buckley P, Howell R (2015) Responses of an ancient woodland field layer to soil translocation: Methods and timing. Applied Vegetation Science 18:579–590

Frostegård Å, Bååth E (1996) The use of phospholipid fatty acid analysis to estimate bacterial and fungal biomass in soil. Biology and Fertility of Soils 22:59–65

Frostegård Å, Tunlid A, Bååth E (1991) Microbial biomass measured as total lipid phosphate in soils of different organic content. Journal of Microbiological Methods 14:151–163

Frostegård Å, Tunlid A, Bååth E (2011) Use and misuse of PLFA measurements in soils. Soil Biology and Biochemistry 43:1621–1625

Hall JE, Kirby KJ, Whitbread AM (2001) National Vegetation Classification: Field guide to woodland. JNCC, Peterborough, UK.

Harris JA (2003) Measurements of the soil microbial community for estimating the success of restoration. European Journal of Soil Science 54:801–808

Harris JA (2009) Soil microbial communities and restoration ecology: facilitators or followers? Science (New York, N.Y.) 325:573–4

Van Der Heijden MGA, Bardgett RD, Van Straalen NM (2008) The unseen majority: soil microbes as drivers of plant diversity and productivity in terrestrial ecosystems. Ecology Letters 11:296–310

Helgason T, Merryweather JW, Young JPW, Fitter AH (2007) Specificity and resilience in the arbuscular mycorrhizal fungi of a natural woodland community. Journal of Ecology 95:623–630

Helliwell DR, Buckley GP, Fordham SJ, Paul TA (1996) Vegetation succession on a relocated ancient woodland soil. Forestry: An International Journal of Forest Research 69:57–74

Hietalahti M, Cadisch G, Buckley GP (2005) The effect of woodland soil translocation on carbon and nitrogen mineralisation processes. Plant and Soil 271:91–107

Hietalahti MK, Buckley GP (2000) The effects of soil translocation on an ancient woodland flora. Aspects of Applied Biology 345–350

Hoeksema JD, Chaudhary VB, Gehring CA, Johnson NC, Karst J, Koide RT, Pringle A, Zabinski C, Bever JD, Moore JC, Wilson GWT, Klironomos JN, Umbanhowar J (2010) A meta-analysis of context-dependency in plant response to inoculation with mycorrhizal fungi. Ecology Letters 13:394–407

Humphries RN (2010) A geomorphological framework & decision tool for habitat translocation practices in the UK. Science 449–467

Jenkinson DS, Powlson DS (1976) The effects of biocidal treatments on metabolism in soil—I. Fumigation with chloroform. Soil Biology and Biochemistry 8:167–177.

Kourtev PS, Ehrenfeld JG, Häggblom M (2003) Experimental analysis of the effect of exotic and native plant species on the structure and function of soil microbial communities. Soil Biology and Biochemistry 35:895–905

MAAF (1986) Ministry of Agriculture Fisheries and Food. The analysis of agricultural materials Reference Book 427. HMSO, London.

Morris R, Alonso I, Jefferson RG, Kirby KJ (2006) The creation of compensatory habitat — Can it secure sustainable development? 14:106–116

Olsson PA (1999) Signature fatty acids provide tools for determination of the distribution and interactions of mycorrhizal fungi in soil. FEMS Microbiology Ecology 29:303–310

Ritz K, Harris JA, Pawlett M, Stone D (2006) Catabolic profiles as an indicator of soil microbial functional diversity. Environment Agency Science Report SC040063/R

Rousk J, Bååth E, Brookes PC, Lauber CL, Lozupone C, Caporaso JG, Knight R, Fierer N (2010) Soil bacterial and fungal communities across a pH gradient in an arable soil. ISME Journal 4:1340–1351

Rousk J, Brookes PC, Bååth E (2010) The microbial PLFA composition as affected by pH in an arable soil. Soil Biology and Biochemistry 42:516–520

Ryan L (2013) Translocation and Ancient Woodland. The Woodland Trust, Grantham, UK.

Smith D (2014) A21 Tonbridge to Pembury Habitat Creation and Enhancement Strategy. WSP UK, Basingstoke, UK.

Wardle DA, Bardgett RD, Klironomos JN, Setälä H, van der Putten WH, Wall DH (2004) Ecological linkages between aboveground and belowground biota. Science (New York, N.Y.) 304:1629–33

Zelles L (1999) Fatty acid patterns of phospholipids and lipopolysaccharides in the characterisation of microbial communities in soil: A review. Biology and Fertility of Soils 29:111–129

Zogg GP, Zak DR, Ringelberg DB, White DC, MacDonald NW, Pregitzer KS (1997) Compositional and Functional Shifts in Microbial Communities Due to Soil Warming. Soil Science Society of America Journal 61:475

